# A phosphorylation switch regulates RAB6 function during mitosis

**DOI:** 10.1101/2023.01.05.522745

**Authors:** Ana Joaquina Jimenez, Hugo Bousquet, Sabine Bardin, Franck Perez, Bruno Goud, Stéphanie Miserey

## Abstract

RAB GTPases are key regulators of membrane trafficking in eukaryotic cells. In addition to their role in interphase, several RAB proteins, including Golgi-associated RAB6, have mitotic functions. The aim of this study was to investigate how the interphasic and mitotic functions of RAB6 could be regulated. Since phosphorylation is a key regulatory process in mitosis, we looked for specific mitotic phosphorylation of RAB6 using a phospho-proteomic approach. We found that RAB6 is phosphorylated at position S52 by the mitotic kinase Pololike kinase 1 (Plk1) in mitosis. Phosphorylated RAB6 localizes at the spindle poles from prophase to anaphase. In metaphase, we observed RAB6A-positive structures containing Mad1 and Mad2 moving along the mitotic spindle via the dynein-dynactin complex. We provide evidence that phosphorylation impairs RAB6A binding to some of its known partners, including p150^*Glued*^ and Bicaudal-D2. In addition, the overexpression of RAB6A phospho-mutants lead to mitosis and cytokinesis defects. Our results suggest that a cycle of RAB6 phosphorylation/dephosphorylation is required for cell division.

## Introduction

Membrane trafficking and mitosis are two essential and interlinked processes in mammalian cells. During mitosis, membrane trafficking along the secretory pathway is downregulated, with the Golgi apparatus disassembling and possibly fusing with the endoplasmic reticulum (ER) (Warren et al., 1983; Featherstone et al., 1985; Zaal et al., 1999). At telophase, the Golgi apparatus reassembles and secretory vesicles are transported to the cleavage furrow for successful cytokinesis (Goss and Toomre, 2008).

In interphase, intracellular membrane trafficking is regulated by RAB GTPases. RAB proteins are localized on the membranes of intracellular compartments, and are specifically involved in the directional movement of transport vesicles from one compartment to another (Zerial and McBride, 2001).

RAB6 is localized to Golgi/TGN (*Trans*-Golgi Network) membranes and plays an essential role in the maintenance of Golgi homeostasis (Goud et al., 2018). There are three main RAB6 isoforms expressed in mammalian cells: RAB6A and RAB6A’ are splice variants of the same gene and are expressed ubiquitously (Echard et al., 2000), while RAB6B is mainly expressed in brain (Opdam et al., 2000). RAB6A’ differs from RAB6A in only three amino acids located in regions flanking the PM3 GTP-binding domain (Echard et al., 2000). Both isoforms interact with all identified RAB6 effectors, except for KIF20A (also known as Rabkinesin-6 and Mklp2) which preferentially interacts with the RAB6A isoform (Echard et al., 2000). RAB6A/A’ co-localize on post-Golgi transport vesicles (Miserey-Lenkei et al., 2010) and regulate several transport pathways at the level of the Golgi complex, including anterograde transport to the plasma membrane, intra-Golgi transport, and Golgi-ER retrograde transport (Goud et al., 2018; Fourriere et al, 2019).

In addition to their role in interphase, several RAB proteins, including RAB1, RAB4, RAB5, RAB11 and RAB35, have mitotic functions (Miserey-Lenkei and Colombo, 2016). Previous studies have shown that RAB6A (Miserey et al., unpublished results) and RAB6A’ participate in the spindle assembly checkpoint (SAC) at the metaphase/anaphase transition (Miserey-Lenkei et al., 2006). This checkpoint senses equal tension at each kinetochore of sister chromatids, and ensures equal partitioning of DNA in the two daughter cells. Two RAB6-binding proteins, p150^*Glued*^, a component of the dynein/dynactin complex which is required for the transport of SAC components (Howell et al., 2001), and GAPCenA, associated to spindle poles in mitosis (Cuif et al., 1999), are both involved in this process (Miserey-Lenkei et al., 2006). Several other RAB6 effectors have also been shown to be involved in mitosis and cytokinesis: KIF20A which plays a key role in the organization of the central spindle at anaphase and cytokinesis (Echard et al., 1998; Hill et al., 2000; Fontijn et al., 2001; Gruneberg et al., 2004); the scaffold protein ELKS, which is part of the cortical microtubule-stabilizing complex (Bouchet et al., 2016; Liu et al., 2016) and the inositol phosphatase OCRL (Dambournet et al., 2011).

The aim of this study was to investigate how RAB6 activity is regulated during cell division. Since phosphorylation is a key regulatory process in mitosis, we looked for specific mitotic phosphorylation of RAB6 using a phospho-proteomic approach. We found that RAB6 is phosphorylated at position S52 by the mitotic kinase Polo-like kinase 1 (Plk1) and that phospho-RAB6 localizes at the spindle poles. We show that RAB6A phosphorylation affects its binding to some of its partners. We provide evidence that disrupting the cycle of RAB6A phosphorylation results in defects in the dynamics of SAC components as well as in cytokinesis.

## Results

### RAB6 is phosphorylated at the spindle poles during mitosis

We performed a phospho-proteomic screen using HeLa cells expressing GFP-RAB6A and GFP-RAB6A’ synchronized in S phase or M phase. Following immunoprecipitation with GFP-Trap, phosphorylation sites were identified using mass spectrometry. We identified five sites differentially phosphorylated on serine residues in interphase and mitosis: three specific to mitosis located at positions 2, 52, and 179 (colored in red in Fig. 1A), one specific to interphase at position 37 (colored in blue in Fig. 1A), and one site phosphorylated in interphase and mitosis at position 117 (colored in green in Fig. 1A). In this study, we focused on the phosphorylation site located at the position S52.

**Figure 1:**
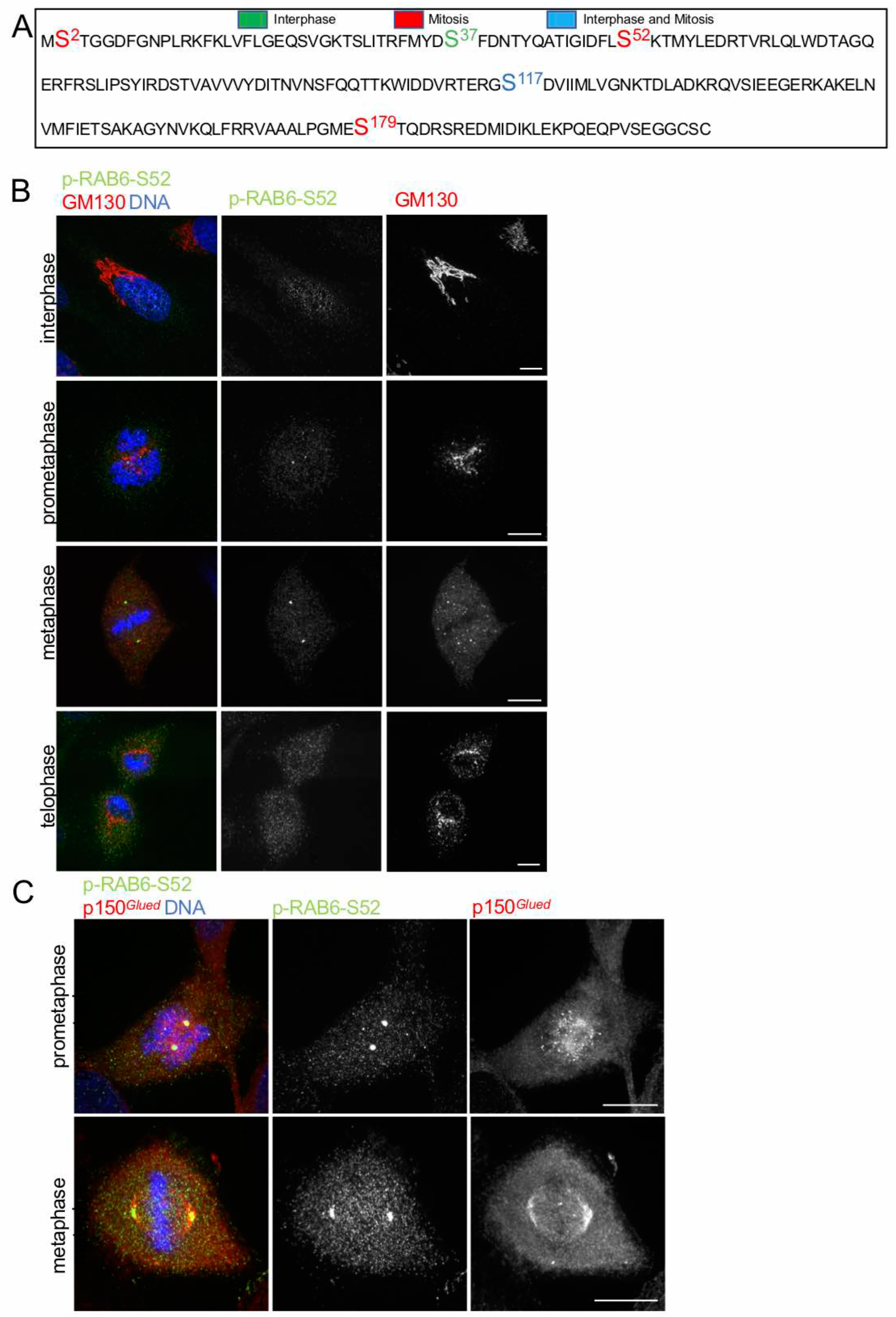
RAB6 is phosphorylated at the spindle poles in mitosis. A/ Amino acid sequence of human RAB6A. The phosphorylation sites specific of interphase (S37), mitosis (S2, S52, 179) and interphase and mitosis (S117) are colored in green, red and blue respectively. B/ Staining of endogenous p-RAB6-S52 (green), GM130 (red) and DNA (blue) in HeLa cells in different phase of the cell cycle: interphase, prometaphase, metaphase, telophase. Bar, 10 μm. C/ Staining of endogenous p-RAB6-S52 (green), p150^*Glued*^ (red) and DNA (blue) in prometaphase and metaphase HeLa cells indicates a colocalization between p-RAB6-S52 and p150^*Glued*^ at the spindle poles. Bar, 10 μm.

We then raised phospho-specific antibodies against a phospho-peptide that includes S52. This antibody is henceforth referred to as p-RAB6-S52. No p-RAB6-S52 staining was observed at the Golgi complex in interphase (Fig. 1B). In mitosis, p-RAB6-S52 staining was observed at two dots reminiscent of the spindle poles (Fig. 1B). To confirm this, cells were co-labelled for p150^*Glued*^, known to be associated with both the kinetochores and the spindle poles in mitosis. The p-RAB6-S52 antibody labeled mainly the spindle poles from prophase to anaphase (Fig. 1C, S1B). A minor pool was observed at kinetochores in prometaphase using BubR1 as a marker (Fig. S1A, arrowheads).

The specificity of p-RAB6-S52 labeling in mitosis was validated by depleting RAB6 using siRNAs and lambda phosphatase treatment (Fig. S1C-D). Aurora B, which localizes to the kinetochores from prophase to metaphase, was used as a marker of mitotic cells. In metaphase cells, p-RAB6-S52 staining at the spindle poles and kinetochores was significantly reduced by 50% following RAB6A or RAB6A’ siRNA treatment and by 80% following RAB6A/A’ siRNA (Fig. S1C). Lambda phosphatase treatment resulted in a total loss of p-RAB6-S52 labeling at the spindle poles (Fig. S1D). These experiments confirmed the specificity of the antibody and indicating that both RAB6A and RAB6A’ are phosphorylated.

### Polo-like kinase 1 phosphorylates RAB6 isoforms during mitosis

To identify the kinase(s) involved in RAB6 phosphorylation in mitosis, cells were treated with different specific inhibitors and the intensity of the p-RAB6-S52 staining at the spindle poles was measured. When cells were treated with staurosporine, a broad-spectrum protein kinase inhibitor, the p-RAB6-S52 signal at the spindle poles labeled with γ-tubulin was markedly reduced in a dose-dependent manner (Fig. S2A). No effect was observed upon treatment with specific CDKs (roscovitine) or MAP kinase (U0126) inhibitors (Fig. S2B). Plk1 is a key mitotic kinase involved in multiple steps of mitosis. Treatment with the specific Plk1 inhibitor BI 2356 at a classical 100 nM concentration leads to cells with a monopolar mitotic spindle (labelled with p150^*Glued*^) (Fig. 2A). In these cells, the labeling of p-RAB6-S52 was completely absent from the spindle poles. The decrease in the intensity of p-RAB6-S52 labeling was dose-dependent (Fig. 2A).

**Figure 2:**
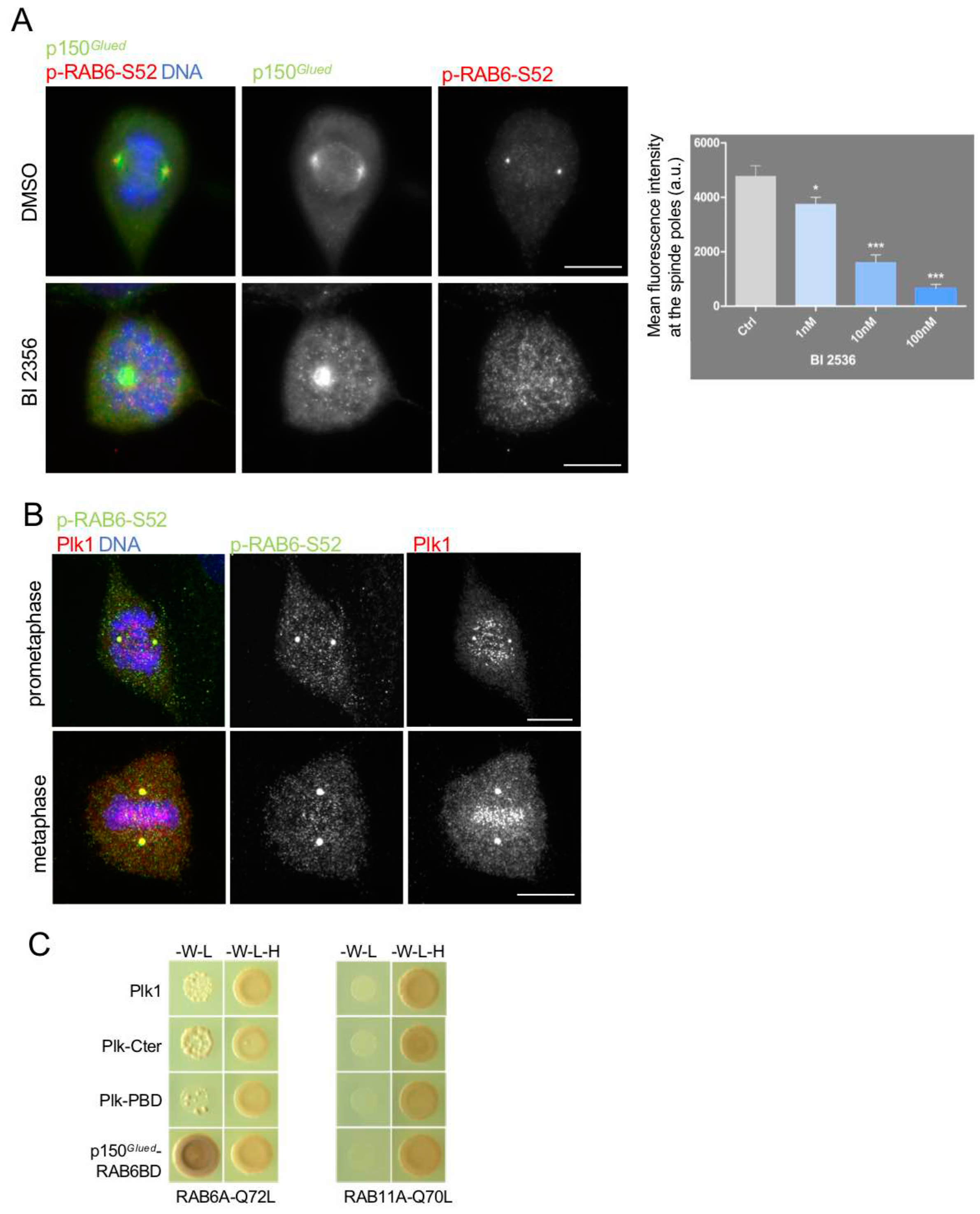
Polo-like kinase phosphorylates RAB6 in mitosis. A/ Left: Staining of endogenous p-RAB6-S52 (red), p150^*Glued*^ (green) and DNA (blue) in HeLa cells treated or not with the Plk1 inhibitor BI 2356. Bar, 10 μm. Right: Quantification of the staining of endogenous p-RAB6-S52 in cells treated as indicated above. (mean ± SEM) * p<0.05, *** p<10^-6^ (Student’s t-test). B/ Staining of endogenous p-RAB6-S52 (green), Plk1 (red) and DNA (blue) in prometaphase and metaphase HeLa cells indicates a colocalization between p-RAB6-S52 and Plk1 at the spindle poles. Bar, 10 μm. C/ Yeast-two hybrid interactions between RAB6A-Q72L or RAB11A-Q72L with different fragments of Plk1. Growth on medium lacking histidine (-W-L-H) indicates an interaction between the encoded proteins.

We then carefully examined the colocalization at the spindle poles between Plk1 and p-RAB6-S52. We found that Plk1 and p-RAB6-S52 colocalize at the spindle poles from pro-metaphase to anaphase, with the highest colocalization observed in metaphase (Fig. 2B). In addition, as mentioned above (Fig. S2B), a pool of p-RAB6-S52 was also found colocalized with Plk1 at kinetochores (data not shown).

We then investigated whether RAB6A could interact with Plk1 (Fig. 2C). Plk1 is composed of a N-terminal Ser/Thr kinase domain, a central interdomain linker, and a C-terminal Polo Box Domain (PBD) involved in its interaction with phosphorylated targets. Using the yeast-two-hybrid assay, we found that the GTP-bound form of RAB6A (RAB6A-Q72L) interacts with Plk1 full length, the C-terminal domain (that contains the PBD domain), and the PBD domain (Fig. 2B). The GTP-bound form of RAB11A was used as a negative control, and the interaction between the RAB6-binding domain of p150^*Glued*^ and RAB6A-Q72L as a positive control.

Our results suggest that p-RAB6-S52 is phosphorylated by Plk1 at the spindle poles in metaphase and is colocalized with Plk1 at the spindle poles and at the kinetochores in prometaphase and metaphase. Additionally, RAB6A directly interacts with the PBD domain of Plk1.

### RAB6 is highly dynamic during mitosis

To investigate the dynamics of RAB6 in mitotic cells, we used time-lapse multicolor videomicroscopy on live cells stably expressing GFP-RAB6A and co-labeled with microtubules (Sir-tubulin) and DNA (Hoechst) (Fig. 3A and Movie S1). In prometaphase and metaphase (Fig. 3A shows an example of a cell in metaphase), we observed highly dynamic GFP-RAB6A-positive structures distributed everywhere in the cytoplasm. To investigate their trajectories, we performed time-projection over a period of five seconds (Fig. 3B). As shown in single images from time-lapse movies, some GFP-RAB6A-positive structures could be seen moving along the mitotic spindle from the kinetochores to the spindle poles (Fig. 3B arrowheads; single time-lapse images are displayed in Fig. S3). Additionally, GFP-RAB6A labelling was also observed associated with the spindle poles (Fig. 3B, arrows).

**Figure 3:**
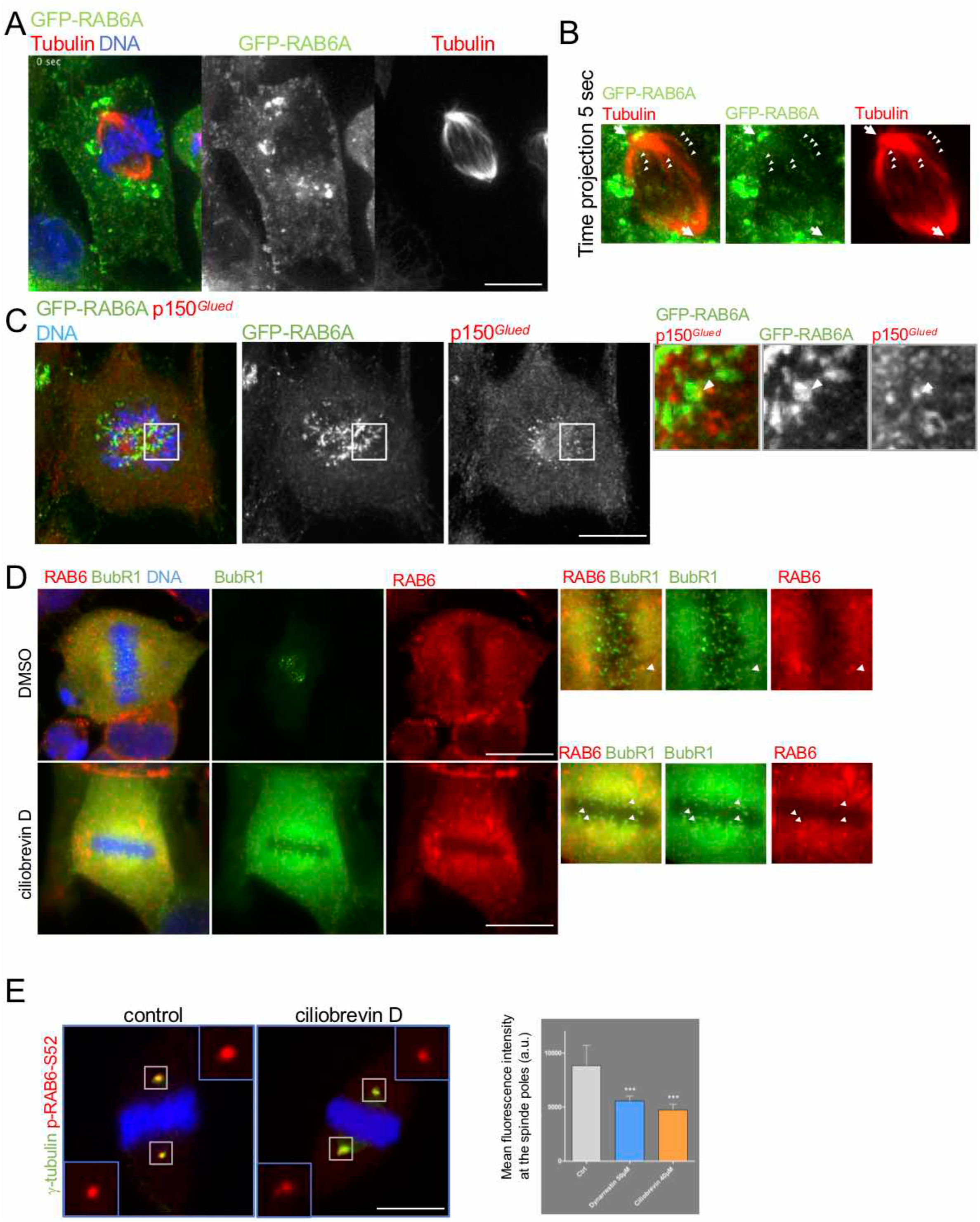
RAB6A is highly dynamic during mitosis. A/ RPE1 cells stably expressing GFP-RAB6A in metaphase were imaged by time-lapse microscopy every second for 30 seconds (see supplementary movie 1). Tubulin is stained in red and DNA in blue. B/ Time projection over 5 seconds of images acquired as described above. Arrowheads indicate GFP-RAB6A structures moving along the mitotic spindle; arrows show GFP-RAB6A positive structures localized near the spindle poles. Bar, 10 μm. C/ Staining of GFP-RAB6A (green), p150^*Glued*^ (red) and DNA (blue) in RPE cells in prometaphase show some structures (arrowheads) where GFP-RAB6A and p150^*Glued*^ colocalize. Bar, 10 μm. D/ Staining of endogenous RAB6 (red), BubR1 (green) and DNA (blue) in HeLa cells treated or not with ciliobrevin D. Arrowheads show RAB6 positive structures colocalizing with BubR1 at kinetochores. Bar, 10 μm. E/ Left: Staining of endogenous p-RAB6-S52 (red), γ-tubulin (green) and DNA (blue) in HeLa cells treated or not with ciliobrevin D. Bar, 10 μm. Right: Quantification of the staining of endogenous p-RAB6-S52 in cells treated as indicated above. (mean ± SEM) *** p<10^-6^ (Student’s t-test).

We then tested the role of the dynein/dynactin complex in the dynamics of GFP-RAB6A positive structures. First, we looked at the co-localization between GFP-RAB6A and p150_*Glued*_ in pro-metaphase and metaphase. Figure 3C shows a cell in pro-metaphase, where some GFP-RAB6A-positive structures are found colocalized with p150^*Glued*^ at kinetochores. Second, to directly test the involvement of the dynein/dynactin complex in the transport of GFP-RAB6A-positive-structures from kinetochores to the spindle poles, cells were treated with the dynein inhibitors ciliobrevin D and dynarrestin, and we measured the amount of endogenous RAB6 at kinetochores (Fig. 3D) and the amount of p-RAB6-S52 labeling at the spindle poles (Fig. 3E). Upon ciliobrevin D treatment, in metaphase cells, the number of kinetochores labeled with endogenous RAB6 was increased by six-fold (BubR1 was used as a marker of kinetochores, Fig. 3D). In addition, following ciliobrevin-D or dynarrestin treatments, the amount of p-RAB6-S52 labeling at the spindle poles, labelled with γ-tubulin, was significantly decreased by 50% (Fig. 3E).

Overall, these results suggest that p150^*Glued*^ and the dynein/dynactin complex are involved in the transport of RAB6-positive structures from kinetochores to the spindle poles along the mitotic spindle. At the spindle poles, the pool of RAB6-positive structures arriving from the kinetochores may then be phosphorylated by Plk1 (see Fig. 6 for a working model).

### RAB6A phosphorylation at Serine 52 affects its binding to some of its partners

The phosphorylation of RAB proteins has been shown to affect their interactions with regulatory proteins (GEFs, GAPs, GDI) and effectors ((Xu et al., 2021). To investigate if it was also the case for RAB6A, we produced a non-phosphorylable RAB6A-S52A mutant and a phospho-mimetic RAB6A-S52D mutant, and assessed their binding to several known RAB6-binding proteins using a yeast two-hybrid assay (Table 1): Bicaudal-D2 (Matanis et al., 2002), p150^*Glued*^, RAB6IP1 (also known as DENND5A)(Miserey-Lenkei et al., 2007), RAB6IP2/ELKS (Grigoriev et al., 2007; Fourriere et al., 2019), KIF20A (Hill et al., 2000; Fontijn et al., 2001), OCRL, GAPCenA (Cuif et al., 1999), and MINT-3 (Teber et al., 2005).

**Table 1.**

|  | Bicaudal-D2-RAB6BD | p150<sup>Glued</sup> RAB6BD | RAB6IP1 (DennD5A) | RABIP2 ELKS | KIF20A RAB6BD | OCRL RAB6BD | GAPCenA RAB6BD | Mint-3 |
| --- | --- | --- | --- | --- | --- | --- | --- | --- |
| RAB6A wt | + | + | + | - | + | - | + | + |
| RAB6A wt-S52A | + | + | n.d | - | + | - | + | n.d |
| RAB6A wt-S52D | - | - | + | - | - | - | + | + |
| RAB6AQ72L | + | + | + | + | + | + | + | + |
| RAB6AQ72L-S52A | n.d. | n.d. | n.d | + | n.d. | + | n.d | n.d |
| RAB6AQ72L-S52D | n.d. | n.d. | + | - | n.d. | - | + | + |

We found that the phosphomimetic RAB6A-S52D mutant was unable to interact with p150^*Glued*^, Bicaudal-D2, ELKS, KIF20A and OCRL whereas the interaction with RAB6IP1, GAPCenA, and MINT-3 was preserved. These results were confirmed by coimmunoprecipitation (data not shown). Conversely, the RAB6A-non-phosphorylable mutant RAB6A-S52A was still able to interact with all tested RAB6-binding proteins (Table 1).

Our results thus suggest that the phosphorylation of RAB6A at Ser 52 impairs its binding to effectors that were shown to play a role in mitosis and cytokinesis.

### RAB6-positive structures contain SAC components

At the metaphase-anaphase transition, the dynein/dynactin complex is directly involved in the transport of components of the SAC from the kinetochores to the spindle poles (Howell et al., 2001). We first verified the colocalization of Mad2 with p150^*Glued*^ at kinetochores in prometaphase. RPE-1 cells expressing Venus-Mad2 were stained with an antibody against p150^*Glued*^. As shown in Fig. S4B, Mad2 fully co-localized with p150^*Glued*^. Mad2 also colocalized with Plk1, which associates with kinetochores in prometaphase (Fig. S4A).

To assess the role of RAB6 in the transport of components of the SAC, we investigated the colocalization of RAB6 with Mad1 and Mad2, in prometaphase and metaphase. As shown in Figure 4A-B, a partial co-localization with Venus-Mad2 and Mad1 could be observed.

**Figure 4:**
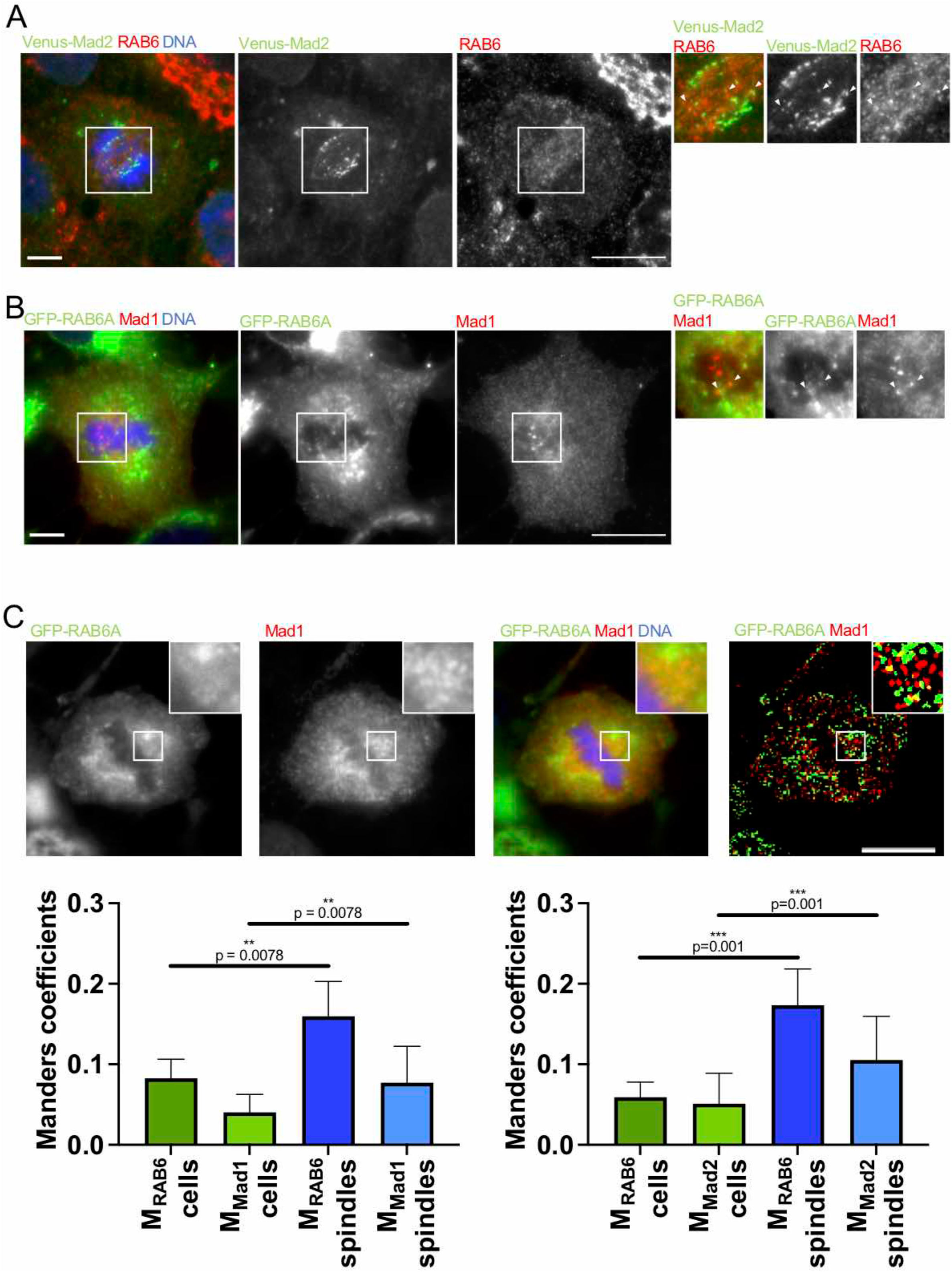
RAB6-positive structures contain components of the spindle assembly checkpoint. A/ Staining of Venus-Mad2 (green), endogenous RAB6 (red) and DNA (blue) in RPE cells in prometaphase show some structures (arrowheads) where Venus-Mad2 and RAB6 co-localize. Bar, 10 μm. B/ Staining of GFP-RAB6A (green), Mad1 (red) and DNA (blue) in HeLa cells show some structures (arrowheads) where GFP-RAB6A and Mad1 co-localize. Bar, 10 μm. C/ Staining of GFP-RAB6A (green), Mad1 (red) and DNA (blue) in HeLa cells. **I**mages were denoised, segmented and binarized in order to determine an intensityindependent coefficient of overlap, based on Manders coefficient calculation. The overlapping coefficients indicate that there is significantly more overlap of both channels in the spindle zone compared to the rest of the cell. Bar, 10 μm.

The colocalization between RAB6, Mad1, and Mad2 at the mitotic spindle in prometaphase and metaphase was further quantified by measuring Manders coefficients (Fig. 4C; see details for quantification in the Material and Methods section). This analysis revealed a significantly higher overlap between RAB6 and Mad1 or Mad2 at the mitotic spindles, kinetochores and spindle poles zone than in the rest of the cell.

These results thus suggest that RAB6 along with the dynein/dynactin complex participates in the transport of the SAC components Mad1 and Mad2.

### Phosphorylation of RAB6A is required for successful mitosis and cytokinesis

We then investigated whether overexpression of RAB6A phosphomutants impacts the cell cycle. HeLa cells expressing GFP-RAB6A-S52A and GFP-RAB6A-S52D were imaged for 72h using time-lapse phase contrast microscopy and their behavior was compared to cells expressing GFP as a control (Fig. 5A). Normal mitosis and successful cytokinesis was observed in GFP expressing cells (Fig. 5A, top, white arrowheads). In contrast, perturbation of cell division was observed in cells overexpressing GFP-RAB6A-S52A. A fraction of the cells initiated mitosis but were unable to complete it for several hours before dying (orange arrowheads, Fig. 5A, bottom), while other cells progressed normally until early cytokinesis but then were unable to separate and dyed (white and blue arrowheads, Fig. 5A, bottom). Quantitative analysis revealed that the expression of GFP-RAB6A-S52A caused a threefold increase in cells dying in mitosis and a seven-fold increase in cells dying during early cytokinesis in comparison to the control situation (Fig. 5B). Intriguingly, the expression of GFP-RAB6A-S52D only led to defective cytokinesis (Fig. 5B).

**Figure 5:**
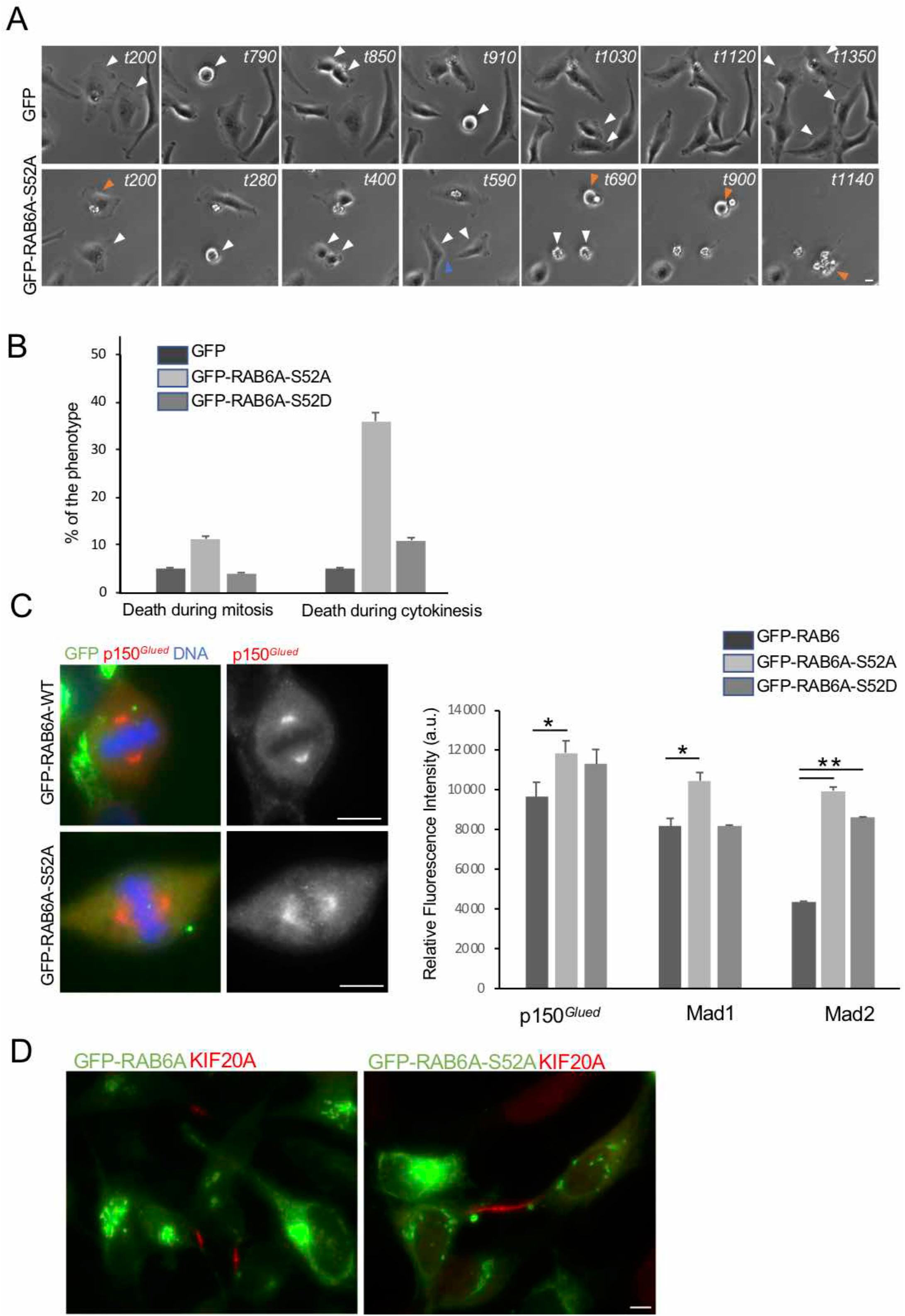
RAB6A phosphorylation is required for successful mitosis and cytokinesis. A/ HeLa cells were transfected with GFP or GFP-RAB6AS52A/D and imaged for 72h by time-lapse videomicroscopy. Top: Arrowheads point to control cells entering mitosis and displaying a successful mitosis and cytokinesis. Bottom: Arrowheads point to GFP-RAB6A-S52A transfected cells entering mitosis and either dying during mitosis (orange) or after cytokinesis (white). Indicated time corresponds to the time in minute after the beginning of the recording. Bar, 10 μm. B/ Quantification of the phenotype observed in cells treated as indicated above. C/ Left: Staining for GFP-RAB6A or GFP-RAB6S52A (green), p150^*Glued*^ (red) and DNA (blue) in HeLa cells. Bar, 10 μm. Right: Quantification of the amount of fluorescent signal corresponding to p150^*Glued*^, Mad1 and Mad2 in cells expressing GFP-RAB6, GFP-RAB6A-S52D or GFP-RAB6A-S52A. (mean ± SEM) * p<0.05, ** p<10^-3^ (Student’s t-test). D/ Staining for GFP-RAB6 or GFP-RAB6S52A (green) and KIF20A (red) in HeLa cells. Bar, 10 μm.

To understand why cells overexpressing the RAB6A phosphomutants die during mitosis, we looked at the localization of components of the SAC and the dynein/dynactin complex. HeLa cells were transfected with GFP-RAB6A-S52A and GFP-RAB6A-S52D and the localization of Mad1, Mad2, and p150^*Glued*^ was observed and quantified in metaphase (Fig. 5C). As shown in Fig. 5C, increased amounts of Mad1, Mad2 and p150^*Glued*^ were found at kinetochores, mitotic spindles, and spindle poles as compared to control cells.

For cells dying in early cytokinesis, we observed that they formed a longer and thicker bridge than in control cells. To quantify their length, we stained the bridges with antibodies against KIF20A (Fig. 5D), Aurora B, and Plk1 (data not shown). A two-fold increase in the length of the bridges was found upon expression of RAB6 phosphomutants.

Altogether, our results suggest that a cycle of phosphorylation/dephosphorylation of RAB6A is critical for successful mitosis and cytokinesis.

## Discussion

In this study, we found that RAB6 is phosphorylated at the spindle poles by Plk1 during mitosis and that its phosphorylation is required for successful mitosis and cytokinesis. That RAB proteins can be phosphorylated has been known since the early 1990s. RAB1 and RAB4 are phosphorylated in mitosis by p34^*cdc2*^ (Cdk1) (Bailly et al., 1991). RAB4 phosphorylation occurs at Ser196, located at the C-terminus region of the protein, and is responsible for the translocation of RAB4 in the cytosol of mitotic cells (van der Sluijs et al., 1992). However, the role of RAB phosphorylation as a regulatory mechanism for their function has gained much attention in recent years since the discovery that the Leucine-rich repeat-2 kinase (LRRK2), whose point mutations are the most common cause of familial Parkinson’s disease (Zimprich et al., 2004; Paisán-Ruíz et al., 2004), phosphorylates a subset of RAB proteins, including RAB3, RAB5, RAB8, RAB10, RAB12, RAB29, RAB35, and RAB43. LRRK2 phosphorylates RAB GTPases at positions 72 and 82 located in the switch 2 region (Jeong et al., 2018; Waschbüsch and Khan, 2020), thus impairing the interaction with effectors. The RAB6 phosphorylation site identified in this study is located at Serine 52 in the interswitch region, a site that has not been previously described for other RAB GTPases. We also identified two additional putative mitotic-specific phosphorylation sites located at S2 and S179, which correspond to consensus sites for Plk1. Preliminary results indicate that the triple phosphomutants give stronger effects on cell cycle progression than the single S52A/D phospho-mutant. One hypothesis is that S2 and S179 act as docking sites for Plk1, enabling Plk1 to phosphorylate RAB6 at S52.

The data presented here suggest that RAB6 plays a phosphorylation-sensitive role in the control of cell division (see Figure 6). We proposed that Mad1, Mad2 and possibly other components of the SAC are transported in RAB6-positive structures moving along microtubules via the dynein-dynactin complex. Upon arrival at the spindle poles, RAB6 is phosphorylated by Plk1, which impairs its interaction with Bicaudal-D2 and p150^*Glued*^, leading to its dissociation from the dynein/dynactin complex. RAB6 could then either be released into the cytosol or re-associates with a kinesin motor to be sent back to the kinetochores. SAC activity does not have on all-or-nothing response and its strength depends on the amount of Mad2 recruited at the kinetochores (Colin et al, 2013). It is thus possible that several RAB6 cycles between kinetochores and the spindle poles take place.

**Figure 6:**
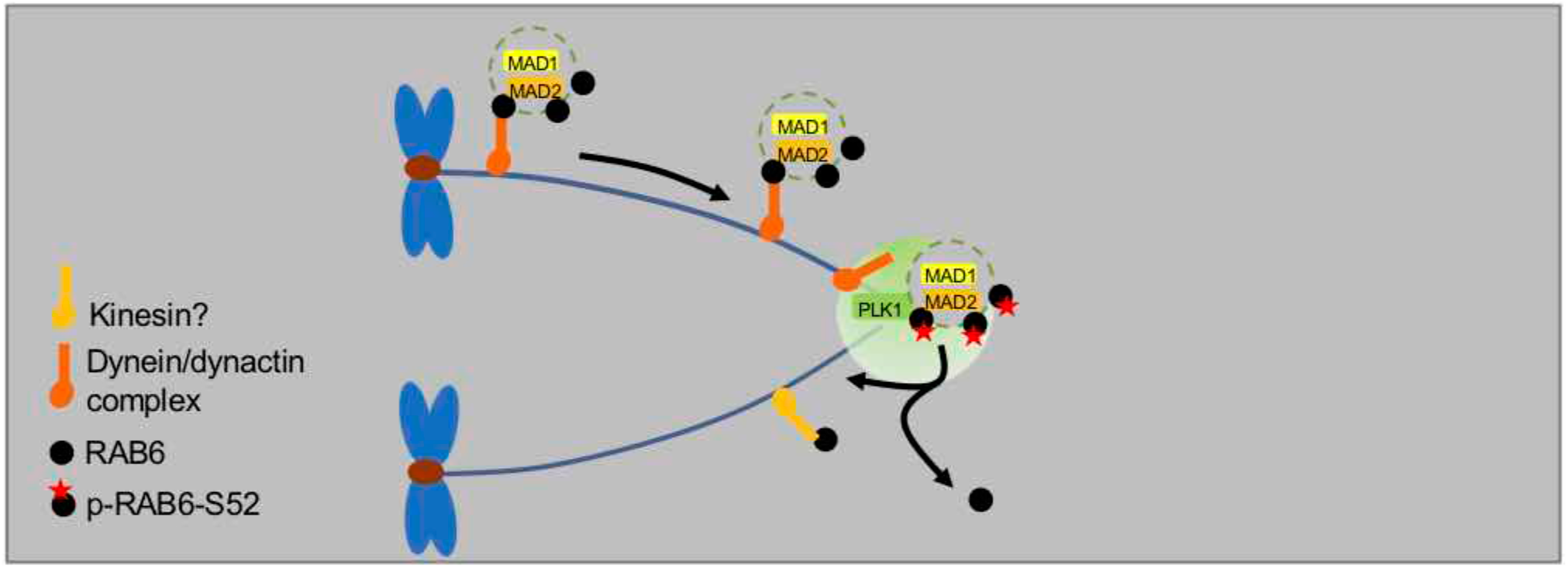
Model for RAB6 function in mitosis. RAB6, along with Mad1 and Mad2, is transported from kinetochores to spindle poles by the dynein-dynactin complex. When RAB6 reaches the spindle poles, it is phosphorylated by Plk1. This phosphorylation prevents the interaction with Bicaudal-D2 and/or p150^*Glued*^, leading to the dissociation of RAB6 from the dynein/dynactin complex. RAB6 could then either be released into the cytosol or re-associated with a kinesin motor to be sent back to the kinetochores.

Several important issues remain to be investigated to validate this model. One is to know whether the RAB6A-positive structures that move along the mitotic spindle correspond to transport vesicles or to protein complexes linked to the dynein-dynactin complex. Of note, we previously obtained evidence that the active pool of RAB6A’ is likely cytosolic in metaphase (Miserey-Lenkei et al., 2006). Another is to determine the relative contribution of RAB6A and RAB6A’ to the mitotic functions of RAB6. S52 is present on both isoforms and silencing of only one isoform does not fully deplete phospho-RAB6 from the spindle poles (Fig. S1C), suggesting that both RAB6A and RAB6A’ are phosphorylated. In addition, our previous work showed that RAB6A’/p150^*Glued*^ is interaction is important for the dynamics of the dynein/dynactin complex at the kinetochores (Miserey-Lenkei et al., 2006). However, RAB6A and RAB6A’ do not have fully redundant functions in interphase (del Nery et al., 2006), this may also be the case in mitosis.

We found that a significant proportion of cells expressing the RAB6 phosphomutants displays cytokinesis defects (Fig. 5). They could correspond to cells that were able to go through the metaphase/anaphase transition but failed to complete cytokinesis. Interestingly, RAB6 phosphorylation impairs its binding to proteins which have been shown to function in cytokinesis, such as ELKS (Liu et al., 2016), OCRL (Dambournet et al., 2011), and KIF20A (Hill et al., 2000; Fontijn et al., 2001). As for the metaphase/anaphase transition, phosphorylation could represent a mechanism to regulate RAB6 function in cytokinesis. Future experiments should address this point.

In summary, our work illustrates that phosphorylation, as documented for other RAB proteins, is a regulatory mechanism for RAB6 function. A tentative hypothesis is that a cycle of phosphorylation/dephosphorylation of RAB6 takes place in mitosis. The identification of the phosphatase (s) involved will be the matter of future experiments.

## Material and Methods

### Antibodies and reagents

The following reagents were used: Ciliobrevin D (Sigma), BI 2356 (SelleckChem), Staurosporine (Tocris), roscovitine (Sigma), Sir-tubulin, Hoechst (Sigma). The following antibodies were used for biochemical experiments: mouse anti-γ-tubulin (Sigma), rabbit anti-γ-tubulin (M. Bornens, Institut Curie), rabbit anti-RAB6 (Goud et al., 1990), mouse anti-p150^*Glued*^ (BD Biosciences), mouse anti-Plk1 (Santa Cruz Biotechnology), mouse anti-BubR1 (BD Biosciences), mouse anti-Mad1 (AbCam), rabbit anti-Mad2 (Sigma), mouse anti-Aurora B (BD Biosciences). The plasmids encoding Plk1 constructs were a kind gift of Thierry Lorca CRBM, Montpellier, France).

### Cell culture and transfection

HeLa and RPE1 cells were grown in DMEM or DMEM/F12 medium (Gibco BRL) supplemented with 10% fetal bovine serum, 100 U/ml penicillin/streptomycin and 2 mM glutamine. Both cell lines were tested free of mycoplasma contamination. Cells were seeded onto 6-well plates, 18- or 12-mm glass coverslips or Fluorodish and grown for 24h before transfection. For expression of the constructs used in this study, HeLa cells were transfected using X-tremeGENE9 (Roche) following the manufacturer’s instructions. For silencing experiments, HeLa cells were transfected with the corresponding RAB6A/A’ siRNA using HiPerFect (Qiagen) following the manufacturer’s instructions.

### RNA interference

The siRNA sequences to deplete human RAB6A/A’ were described in (del Nery et al., 2006). These siRNAs and the one targeting luciferase (CGUACGCGGAAUACUUCGA) were obtained from Sigma.

### Yeast two-hybrid experiments

Yeast two hybrid experiments were performed as described in (Miserey-Lenkei et al., 2010).

### Construction of RAB6 phospho-mutants

Single point mutations were inserted in the human *RAB6A* sequence at position 156 using the QuickChange mutagenesis kit (Agilent).

### Co-immunoprecipitation and proteomic analysis

HeLa cells transfected with GFP or GFP-RAB6 constructs were treated or not with either 10 μM nocodazole overnight (to block cells in M phase) or 20 μM hydroxyurea (to block cells in S phase), trypsinized, washed once in PBS and incubated on ice for 60 min in a lysis buffer: 25 mM Tris pH 7.5, 50 mM NaCl and 0.1% NP40. Cells were then centrifuged 10 min at 10,000g to remove cell debris. Extracts were then processed for coimmunoprecipitation using GFP-trap (Chromotek) following manufacturer’s instructions. Mass spectrometry analysis was performed at the Proteomic platform of Institut Jacques Monod (Paris, France).

### Immunofluorescence microscopy

HeLa (Echard et al., 1998), RPE1-GFP-RAB6A (Schauer et al., 2010) or RPE-Venus-Mad2 (Collin et al., 2013) cells grown on coverslips were fixed in 4% paraformaldehyde (PFA) for 15 min at RT or in methanol (2 min, −20°C) Cells were then processed for immunofluorescence as previously described (del Nery et al., 2006). The following primary antibodies were used: mouse anti-GM130 (1:1000), mouse anti-γ-tubulin (1:200), rabbit anti-γ–tubulin (1:400), rabbit anti-RAB6 (1:2000), mouse anti-p150^*Glued*^ (1:200), mouse anti-Plk1 (1:200), mouse anti-BubR1 (1:200), mouse anti-Mad1 (1:200), rabbit anti-Mad2 (1:200), mouse anti-Aurora B (1:200). Fluorescently-coupled secondary antibodies were obtained from Jackson. Coverslips were mounted in Mowiol and examined under a 3D deconvolution microscope (Leica DM-RXA2), equipped with a piezo z-drive (Physik Instrument) and a 100×1.4NA-PL-APO objective lens for optical sectioning or spinning-disk microscope mounted on an inverted motorized microscope (Nikon TE2000-U) through a 100×1.4NA PL-APO objective lens. 3D or 1D multicolor image stacks were acquired using the Metamorph software (MDS) through a cooled CCD-camera (Photometrics Coolsnap HQ).

### Time lapse fluorescence microscopy

Transfected cells were grown either on glass bottom Fluorodish or on glass coverslips transferred, just before observation, to custom-built microscope slide chambers (Chamlid chamber). Time-lapse imaging was performed at 37°C using a spinning-disk microscope mounted on an inverted motorized microscope (Nikon TE2000-U) through a 100×1.4NA PL-APO objective lens. The apparatus is composed of a Yokogawa CSU-22 spinning disk head, a Roper Scientific laser launch, a Photometrics Coolsnap HQ2 CCD camera for image acquisition and Metamorph software (MDS) to control the setup. Acquisition parameters were 100 msec exposure for GFP channel and 100 msec for mCherry channel. Laser was set to 30% in each case. Images shown in figures correspond to the maximal intensity projection through the Z axis performed with the Image J software (NIH Image).

### Co-localization quantification

Images were analyzed with custom semi-automated macros using ImageJ (Schneider et al., 2012). Image stacks were denoised by subtraction of mean-blurred self (radius 15). Small objects were then enhanced by subtraction of gaussian-blurred self (radius 2). The resulting image was smoothened by applying a gaussian-blur (radius 1). Each image was thresholded manually to remove extra background, to adapt the best possible to the signal of each image, and subsequently converted into a binary mask. Regions of interest delimiting the spindle area and the surrounding cytoplasm deprived from the spindle area were also determined manually. Binary masks were the applied to original images to calculate the thresholded Manders’ coefficients using the formulas (Manders et al., 1993; Cordelières and Bolte, 2014):

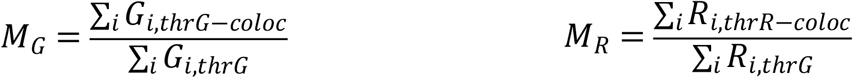

where M_G_ and M_R_ represent Manders’ thresholded coefficient for the green and red channels respectively. As an example, the numerator of the M_G_ equation is the sum of the intensities in the green image of all pixels that are non-null in both channels after thresholding, and within the considered region of interest. The denominator is the sum of all non-null pixels in the green channel and within the region of interest.

### Statistical analysis

All data were generated from cells pooled from at least 3 independent experiments represented as (*n*), unless mentioned, in corresponding legends. Statistical data were presented as means ± standard error of the mean (S.E.M.). Statistical significance was determined by Student’s t-test for two or three sets of data using Excel, no sample was excluded, except for Fig. 4C where Wilcoxon matched pairs rank test was used. Between 24 to 74 cells were quantified. Cells were randomly selected. Only *P*-value <0.05 was considered as statistically significant.

## Supplemental Information

Supplemental Information comprises 4 Supplementary Figures, 1 Supplementary Movie, 1 Supplementary Table, associated legends and references.

## Acknowledgments

We are grateful to Michel Bornens, Ana-Maria Lennon-Duménil and Yohanns Bellaïche for insightful discussions. The authors greatly acknowledge the Nikon Imaging Center @ Institut Curie-CNRS, the PICT-IBiSA, member of the France-BioImaging national research infrastructure. This work was supported by an ERC (European Research Council) advanced grant (project 339847 ‘MYODYN’), an ARC grant (SL220120605302) and a “La Ligue contre le cancer” grant (RS22/75-64). The Goud team is member of Labex CelTisPhyBio (11-LBX-0038), Idex Paris Sciences et Lettres (ANR-10-IDEX-0001-02 PSL) and the “Institut de convergence” Qlife.

## Figure legends

**Figure S1:**
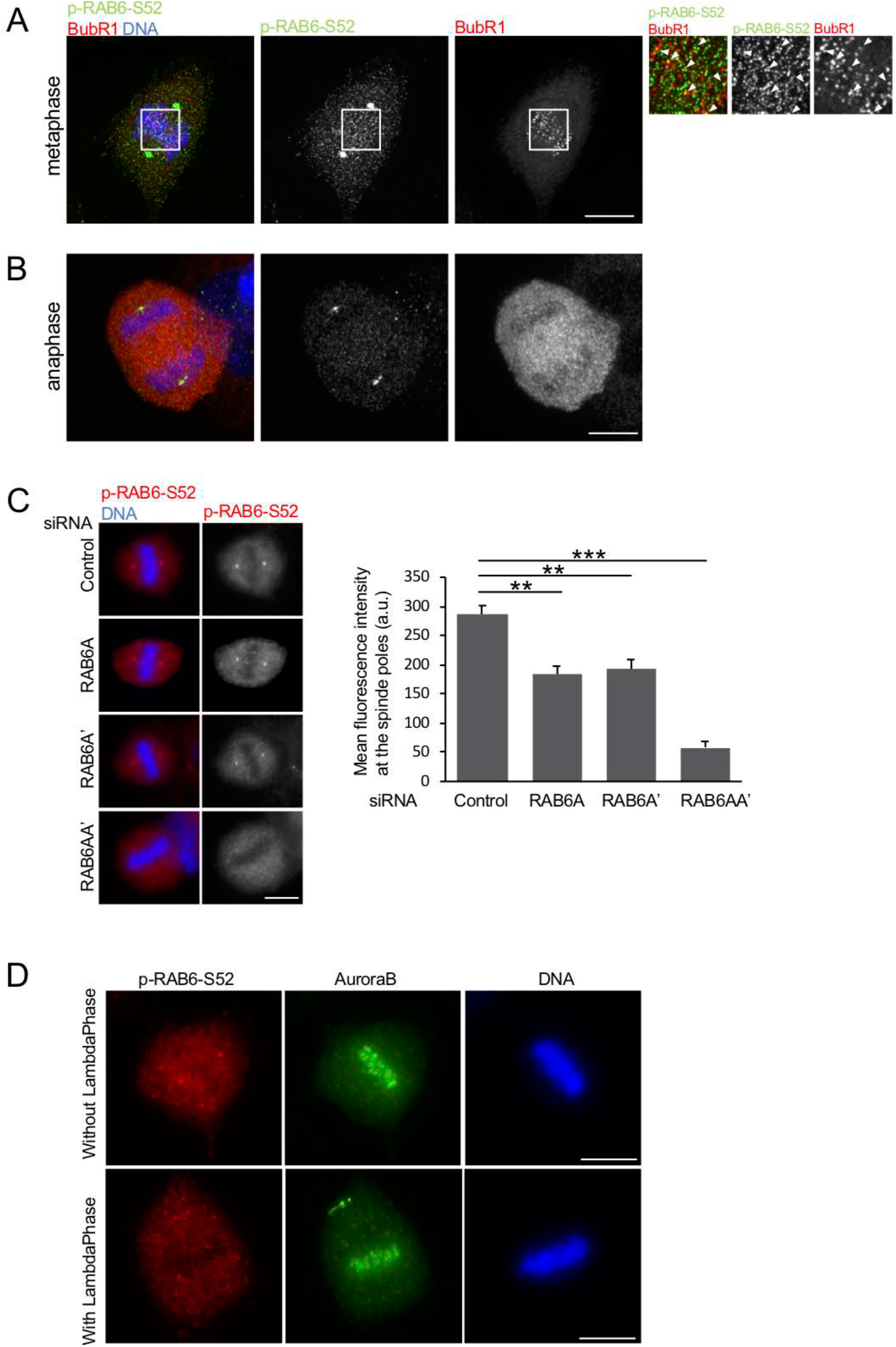
RAB6 is phosphorylated at the spindle poles in mitosis. A/ Staining of endogenous p-RAB6-S52 (green), BubR1 (red) and DNA (blue) in cells in metaphase indicates a colocalization between p-RAB6-S52 and BubR1 at some kinetochores. Bar, 10 μm. B/ Staining of endogenous p-RAB6-S52 (green), BubR1 (red) and DNA (blue) in cells in anaphase indicates a localization of p-RAB6-S52 at the spindle poles. Bar, 10 μm. C/ Left: Staining of endogenous p-RAB6-S52 (red) and DNA (blue) in HeLa cells 3 days after transfection with RAB6A, RAB6A’ or RAB6A/A’ siRNAs. Bar, 10 μm. Right: Quantification of the staining of endogenous p-RAB6-S52 in cells treated as indicated above (mean ± SEM) ** p<10^-4^, *** p<10^-11^ (Student’s t-test) D/ Staining of endogenous p-RAB6-S52 (red), Aurora B (green) and DNA (blue) in HeLa cells treated or not with Lambda Phosphatase. Bar, 10 μm.

**Figure S2:**
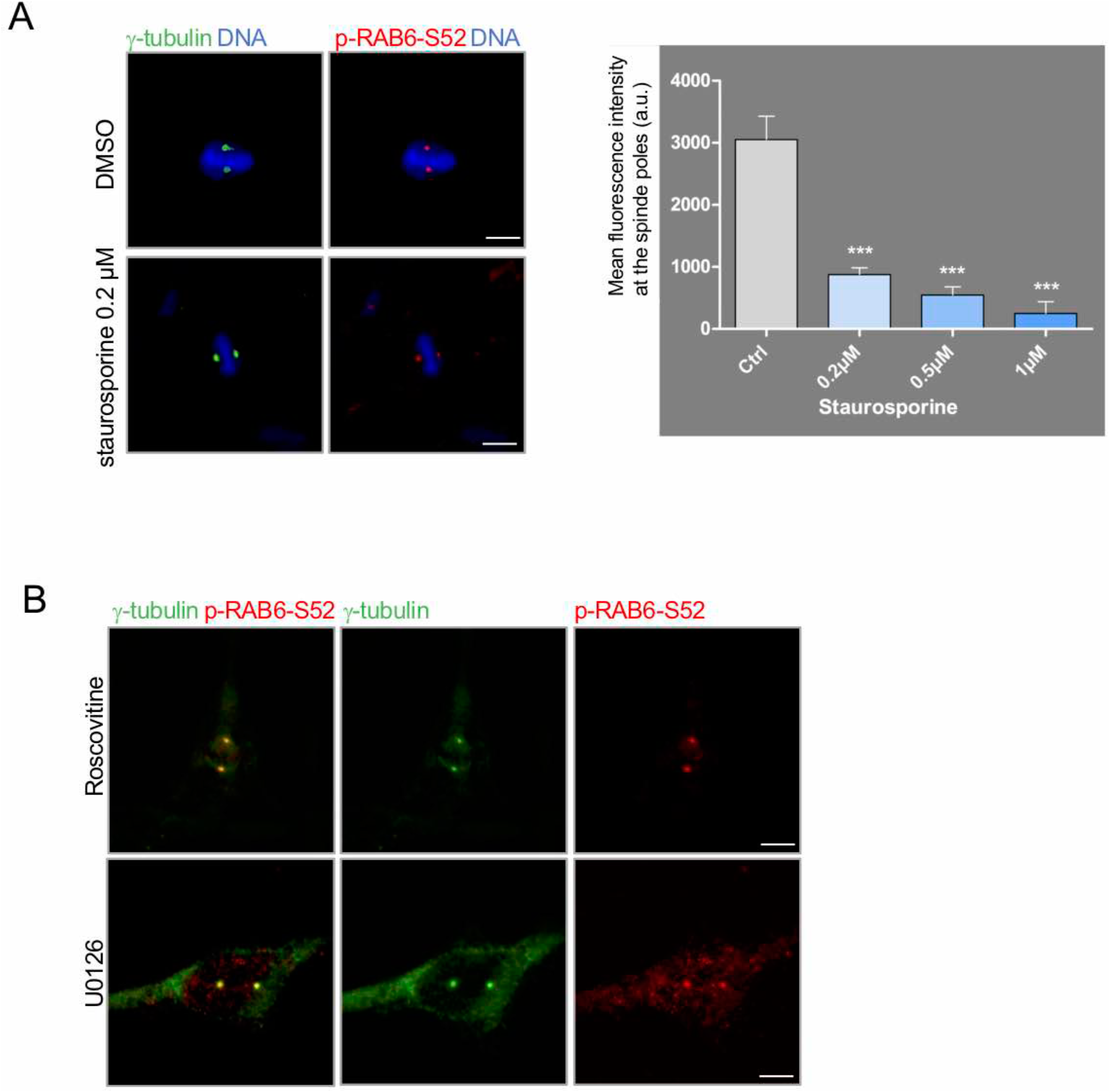
Polo-like kinase phosphorylates RAB6 in mitosis. A/ Left: Staining of endogenous p-RAB6-S52 (red), γ-tubulin (green) and DNA (blue) in HeLa cells treated or not with staurosporine. Bar, 10 μm. Right: Quantification of the staining of endogenous p-RAB6-S52 in cells treated as indicated above. (mean ± SEM) *** p<10^-6^ (Student’s t-test). B/ Staining of endogenous p-RAB6-S52 (red), γ-tubulin (green) and DNA (blue) in HeLa cells treated or not with U0126 or roscovitine. Bar, 10 μm.

**Figure S3:**
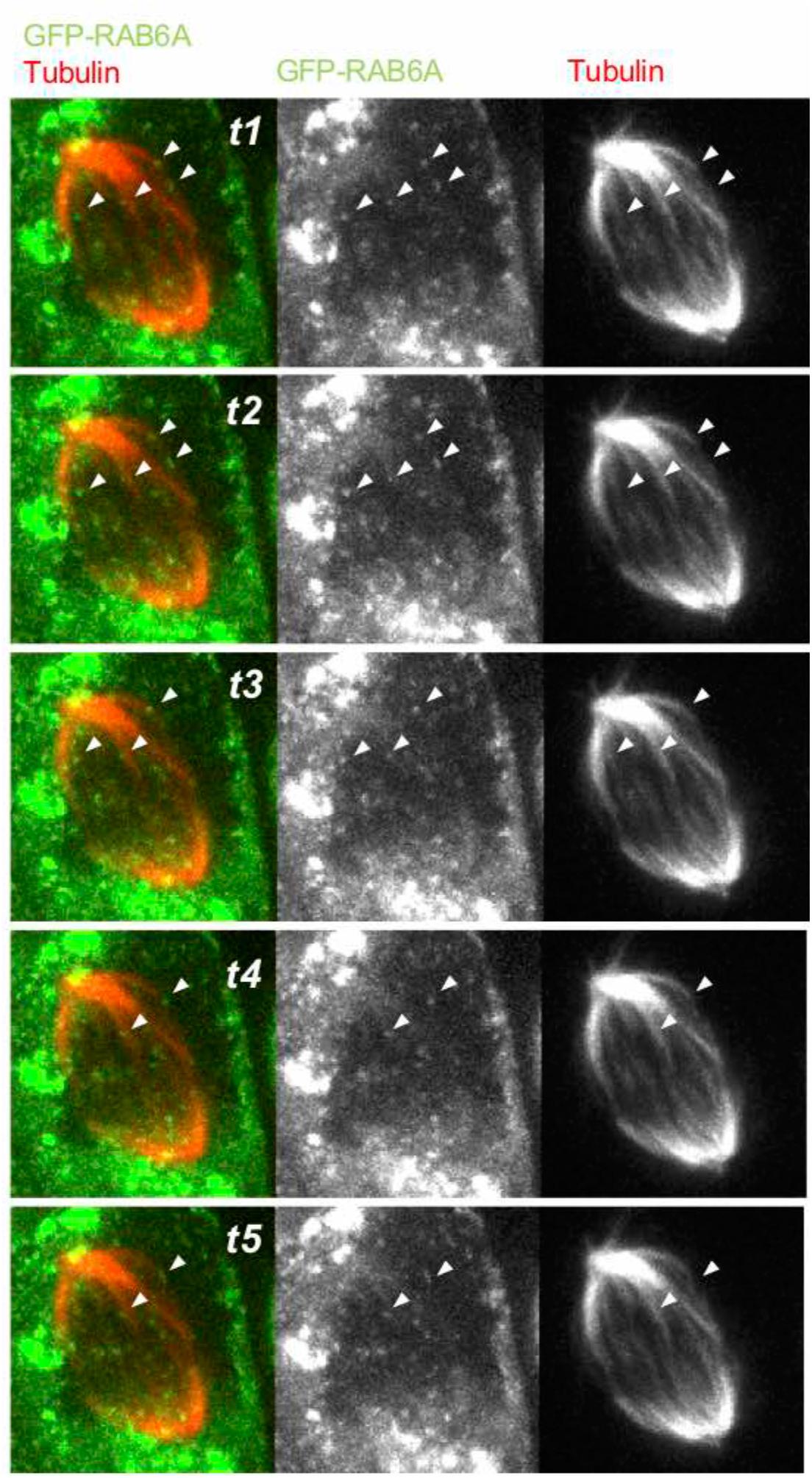
RAB6 is highly dynamic during mitosis. A/ Images from time point 1 sec to time point 5 sec from movies displayed in Figure 3A. Tubulin is stained in red and DNA in blue. Arrowheads indicate GFP-RAB6 structures moving along the mitotic spindle

**Figure S4:**
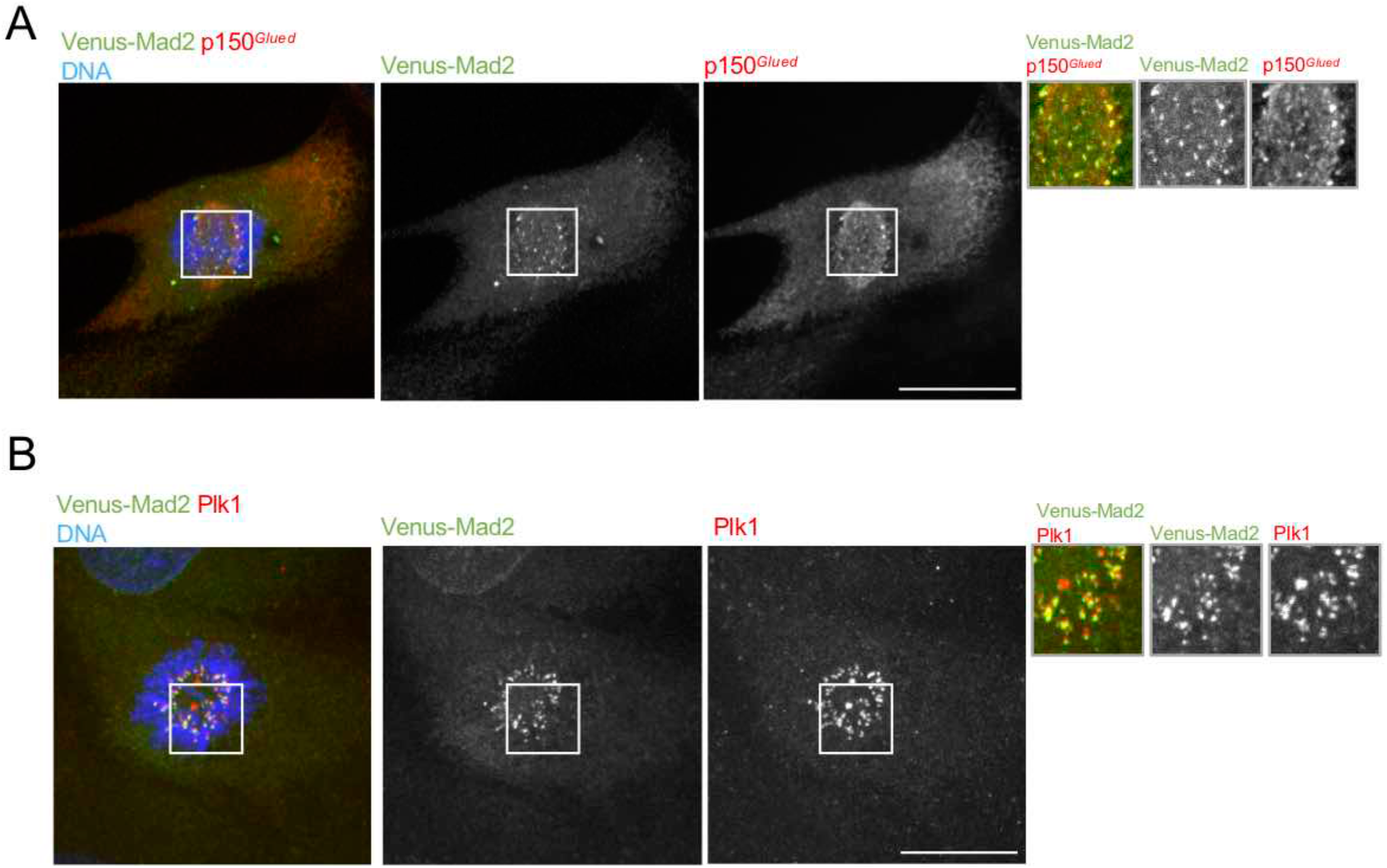
RAB6 transports components of the Mad2 spindle checkpoint. A/ Staining of Venus-Mad2 (green), Plk1 (red) and DNA (blue) in RPE cells in prometaphase shows colocalization between Venus-Mad2 and Plk1. Bar, 10 μm. B/ Staining of Venus-Mad2 (green), p150^*Glued*^ (red) and DNA (blue) in RPE cells in prometaphase shows colocalization between Venus-Mad2 and p150^*Glued*^. Bar, 10 μm.

